# Bumblebee worker castes show differences in allele-specific DNA methylation and allele-specific expression

**DOI:** 10.1101/2020.02.07.938423

**Authors:** H. Marshall, A.R.C. Jones, Z.N. Lonsdale, E.B. Mallon

**Affiliations:** Department of Genetics and Genome Biology, The University of Leicester, Leicester, UK

**Keywords:** genomic imprinting, bumblebee, Hymenoptera, parent-of-origin

## Abstract

Allele-specific expression is when one allele of a gene shows higher levels of expression compared to the other allele, in a diploid organism. Genomic imprinting is an extreme example of this, where some genes exhibit allele-specific expression in a parent-of-origin manner. Recent work has identified potentially imprinted genes in species of Hymenoptera. However, the molecular mechanism which drives this allelic expression bias remains unknown. In mammals DNA methylation is often associated with imprinted genes. DNA methylation systems have been described in species of Hymenoptera, providing a candidate imprinting mechanism. Using previously generated RNA-Seq and whole genome bisulfite sequencing from reproductive and sterile bumblebee (Bombus terrestris) workers we have identified genome-wide allele-specific expression and allele-specific DNA methylation. The majority of genes displaying allele-specific expression are common between reproductive castes and the proportion of allele-specific expression bias generally varies between colonies. We have also identified genome-wide allele-specific DNA methylation patterns in both castes. There is no significant overlap between genes showing allele-specific expression and allele-specific methylation. These results indicate that DNA methylation does not directly drive genome-wide allele-specific expression in this species. Only a small number of the genes identified may be ‘imprinted’ and it may be these genes which are associated with allele-specific DNA methylation. Future work utilising reciprocal crosses to identify parent-of-origin DNA methylation will further clarify the role of DNA methylation in parent-of-origin allele-specific expression.

## Introduction

Allele-specific expression is when one allele of a gene shows higher levels of expression compared to the other allele in a diploid organism. It has been associated with genomic mechanisms such as X-chromosome inactivation and genomic imprinting, i.e. parent-of-origin allele-specific expression (Knight, 2004). It has been predicted that social insects should display imprinted genes (Queller, 2003) based on assumptions of the kinship theory (Haig, 2000). Recent research has identified parent-of-origin allele-specific expression in honeybees and bumblebees (Galbraith *et al.*, 2016; Kocher *et al.*, 2015; Marshall *et al.*, 2020), with one study identifying greater paternal-allele (patrigene) expression bias in reproductive honeybee workers compared to sterile workers, as predicted by the kinship theory (Galbraith *et al.*, 2016). However, the mechanism by which these genes exhibit this expression bias remains unknown.

In mammals and angiosperm plants imprinted genes are often associated with allele-specific DNA methylation (Barlow and Bartolomei, 2014). Many social insects have functional DNA methylation systems, including the eusocial honeybee (Bewick *et al.*, 2016; Lyko *et al.*, 2010) and primitively eusocial bumblebee, *Bombus terrestris* (Sadd *et al.*, 2015). However, the function of DNA methylation in insects remains debated (Glastad *et al.*, 2018).

Various studies have found an association between methylation and gene expression (Glastad *et al.*, 2014; Bonasio *et al.*, 2012; Patalano *et al.*, 2015; Marshall *et al.*, 2019), and alternative splicing (Lyko *et al.*, 2010; Glastad *et al.*, 2016) in social insects. However, this is not uniform across all species, see Standage *et al.* (2016). Additionally, allele-specific expression has been associated with allele-specific methylation in two ant species, *Camponotus floridanus* and *Harpegnathos saltator* (Bonasio *et al.*, 2012). Another study did not find any relationship between allele-specific expression and methylation in a hybrid cross of two non-social wasp species, *Nasonia vitripennis* and *Nasonia giraulti* (Wang *et al.*, 2016).

Bumblebees provide an ideal system to further investigate the relationship between allele-specific methylation and allele-specific expression, specifically with a view of elucidating potential mechanisms involved in genomic imprinting in social insects. Using a candidate gene approach, previous research identified allele-specific expression in a gene (ecdysone 20-monooxygenase-like) related to worker reproductive behaviour in *B. terrestis* (Amarasinghe *et al.*, 2015). Additional research has since used RNA-seq data to identify >500 loci showing allele-specific expression throughout the *B. terrestris* genome (Lonsdale *et al.*, 2017). This same study also identified 19 genes displaying allele-specific expression and allele-specific methylation, although this was in a single individual (Lonsdale *et al.*, 2017).

It is predicted that imprinted genes in *B. terrestris* will specifically have a role regarding worker reproductive behaviour (Queller, 2003). Methylation has been directly associated with reproductive behaviour in *B. terrestris* (Amarasinghe *et al.*, 2014) and recent research has identified differentially methylated genes between reproductive and sterile worker castes (Marshall *et al.*, 2019)

In order to identify the genome-wide relationship between allele-specific expression and allele-specific methylation in *B. terrestris* we have taken advantaged of a previously generated data set. These data consist of whole genome bisulfite sequencing and RNA-seq from reproductive and sterile workers, spanning three genetically distinct colonies. We hypothesise that if genomic imprinting plays a role in worker reproductive behaviour in *B. terrestris*, genes showing allele-specific expression will be enriched for reproductive processes. Additionally, if DNA methylation acts as an imprinting mark in *B. terrestris* then we predict that some genes will show a direct association between allele-specific expression and allele-specific methylation.

## Materials and Methods

### Samples and data

The data used in this study were generated in previously published work by Marshall *et al.* (2019). Briefly, these consist of 18 RNA-Seq libraries generated from head tissue of three reproductive workers and three sterile workers per colony, with three independent colonies total. DNA from head tissue from the same individuals was pooled by reproductive status and colony for whole genome bisulfite sequencing, producing one representative reproductive sample and one sterile sample per colony replicate, giving six whole genome bisulfite libraries total. One RNA-Seq sample, J8_24, was excluded from this study as it was possibly incorrectly labelled in the previous work, see Marshall *et al.* (2019).

### Identification of allele-specific expression

Data were quality checked using fastqc v.0.11.5 (Andrews, 2010) and trimmed using CutAdapt v1.1 (Martin, 2011). Trimmed data were aligned to the reference genome (Bter_1.0, Refseq accession no. GCF_000214255.1 (Sadd *et al.*, 2015)) using STAR v2.5.2 (Dobin *et al.*, 2016) with standard parameters. SNPs were the calling following the GATK best practices for SNP calling from RNA-Seq data (Auwera, 2014). Briefly this involves assigning read groups and marking duplicate reads using Picard v.2.6.0 (Broad Institute, 2018), removing reads overlapping introns to keep only exonic reads, calling SNPs with a minimum confidence score of 20.0, then filtering SNPs by windows of three within a 35bp region, to keep only those with a Fisher strand value greater than 30.0 and a quality by depth value greater than 2.0 (these filtering steps are considered particularly stringent) (Auwera, 2014). These SNPs were then incorporated into the WASP v.0.3.1 pipeline (van de Geijn *et al.*, 2015) which re-maps all reads with either the reference SNP or alternative SNP in order to reduce reference allele mapping bias. Reads that cannot be mapped with the alternative SNP are discarded. SNPs were then filtered to keep only biallelic SNPs allowing individual alleles to be identified. Final reads were then counted per biallelic SNP using the ‘ASEreadcounter’ program from GATK.

A custom R script was used to annotate the SNP positions with gene identifiers, SNPs were filtered to remove those with a coverage of less than 10. SNPs were also removed if they had a count of zero for either the alternative or reference SNP as they may have been mis-called by the SNP caller as heterozygous when they are actually homozygous. Two new columns were then created to represent each allele, as it is not possible to tell which SNPs belong to which allele (e.g. a reference SNP at a given position may be accompanied with an alternative SNP on the same allele). The counts for each SNP were then allocated to either ‘allele: 1’ or ‘allele: 2’, with the highest counts per SNP allocated to ‘allele: 1’ (Fig.1 and supplementary 2.0 Fig.S1). Counts per SNP per allele were then summed over each gene for each reproductive status per colony creating one representative sample per reproductive status per colony. Conducting analyses on a per gene basis decreases false positive calls of allele-specific expression which may occur if there is some remaining reference allele mapping bias after re-mapping with WASP (Degner *et al.*, 2009).

**Figure 1:**
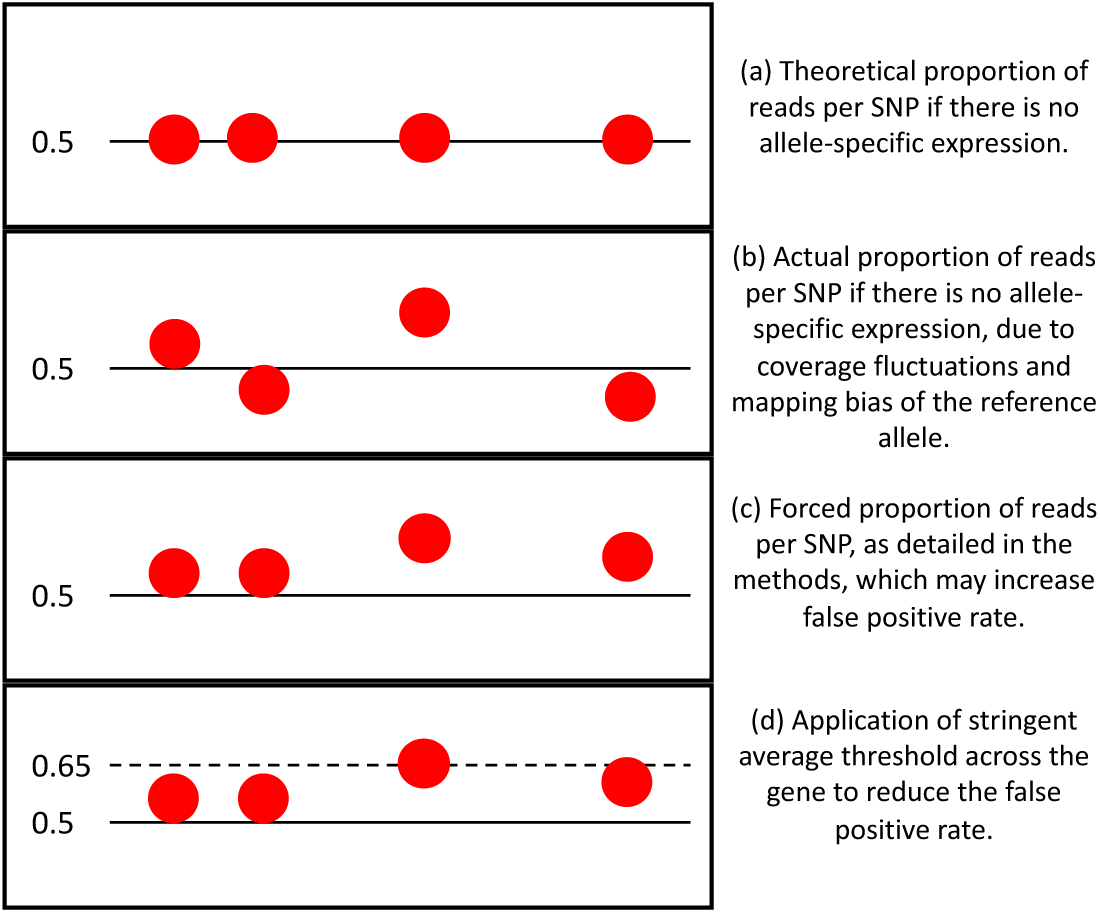
Overview of the theoretical proportions of reads per SNP in a gene which does not show allele-specific expression. Each red dot is an individual SNP.

As this method is naive to allele specific alternative splicing, stringent filtering was applied throughout. Only genes with counts found in at least two of the three colony replicates per reproductive caste were tested. A logistic regression model was then applied with the proportion of allelic expression per gene as the dependent variable and with reproductive status and colony as independent variables, a quasibiomial distribution was applied to account for any overdispersion within the data. P-values were corrected for multiple testing using the Benjimini-Hochberg method (Benjamini and Hochberg, 1995) and genes were classed as showing allele specific expression if the q-value was <0.05 and the average proportion of allelic expression per caste across colonies was >0.65. This stringent filtering was used to account for cases of mis-allocation of SNPs to the correct alleles (Fig.1).

### Identification of allele-specific methylation

Data quality were checked using fastqc v.0.11.5 (Andrews, 2010) and trimmed using CutAdapt v1.1 (Martin, 2011). Trimmed data were aligned to the reference genome (Bter_1.0, Refseq accession no. GCF_000214255.1, (Sadd *et al.*, 2015)) using Bismark v.0.16.1 (Krueger and Andrews, 2011) and bowtie2 v.2.2.6 (Langmead and Salzberg, 2012) with standard parameters. Alignment output files were deduplicated using Bismark v.0.16.1 (Krueger and Andrews, 2011) and sorted and indexed using samtools v.1.3.2 (Li *et al.*, 2009).

Allele-specific methylation was determined using a probabilistic model implemented using the ‘*amrfinder*’ program from the MethPipe package v.3.4.2 (Fang *et al.*, 2012). This program scans the genome using a sliding window approach and fits two models to each interval, one model predicts the methylation levels of each window are the same for both alleles and a second model predicts the methylation levels are different for each allele. The likelihood of the two models is then compared and a false discovery rate corrected p-value is generated per window (Fang *et al.*, 2012). Sample input files were merged by reproductive group in order to increase the coverage per CpG as this method does not take replication into account. Windows were defined as three CpGs with a minimum coverage of 10 reads per CpG. Only regions within the main 18 linkage groups of the *B. terrestris* genome were tested for allele specific methylation as the program is not designed to cope with the number of unplaced scaffolds (5,591) that the current genome build contains. Finally, allelically methylated regions falling within a gene were annotated with the gene identifier using a custom R script.

This method of identifying allelically methylated regions is preferable compared to using SNP data to identify alleles for the data presented here. Firstly, it is difficult to call SNPs reliably from bisulfite data, this is because C/T SNPs and C/T conversions introduced during bisulfite treatment appear the same within the data (Liu *et al.*, 2012). Secondly, as the samples used were pooled females, each sample may contain multiple SNPs at a given loci meaning the coverage produced per SNP would be too low to produce any reliable estimates of allelic methylation.

### Gene ontology analysis

Gene ontology terms for *B. terrestris* were taken from a custom database made in Bebane *et al.* (2019). GO enrichment analysis was carried out using the hypergeometric test with Benjamini-Hochberg (Benjamini and Hochberg, 1995) multiple-testing correction, q <0.05. GO terms from genes showing allele-specific expression were tested for enrichment against a database made from the GO terms of all genes identified in the RNA-Seq data. GO terms from genes showing allele-specific methylation were tested for enrichment against a database made from the GO terms of all genes identified as methylated. Genes were determined as methylated if they had a mean weighted methylation level (Schultz *et al.*, 2012) greater than the bisulfite conversion error rate of >0.05. Descriptions of GO terms and treemaps were generated by REVIGO (Supek *et al.*, 2011).

### Relationship between allele-specific expression and allele-specific methylation

Significant overlap between genes showing allele-specific expression and allele-specific methylation was tested using a hypergeometric test. Overlap plots were generated using the *UpSetR* package in R (Lex *et al.*, 2016). Custom R scripts were used to test for a relationship between allele-specific expression and allelically methylated genes and the interaction of that relationship with reproductive caste.

## Results

### Allele-specific expression

All reads had 13bp trimmed from the start due to base bias generated by the Illumina protocol (Krueger *et al.*, 2011). The mean number of uniquely mapped reads was 89.4% ± 0.8% (mean ± standard deviation). This equated to a mean of 10,115,366 ± 1,849,600 uniquely mapped reads (supplementary 1.0.0). The average number of heterozygous SNPs called per sample was 17,753 ± 6,840, of which an average of 9,355 ± 3,781 had a coverage greater than 10 and after filtering to remove potentially homozygous SNPs the average final number of SNPs per sample was 9,297 ± 3,755 (supplementary 2.0, Fig.S2a). The average number of genes with at least one SNP per sample was 2,436 ± 947 (supplementary 2.0, Fig.S2b).

Only genes present in at least two colonies per reproductive status were tested for allele-specific expression, this lead to a final conservative list of 2,673 genes (24.2% of all annotated genes in the reference genome Bter_1.0). A total of 139 genes were found to show significant allelic expression bias (q <0.05 and average allelic expression proportion >0.65), supplementary 1.0.1 and 2.0, Fig.S3. As expected there were many genes which show a significant q-value below the cut-off threshold of 0.65 (supplementary 2.0, Fig.S4).

The genes of reproductive and sterile workers show similar levels of allelic expression (Spearman’s rank correlation, S = 1229363078, rho = 0.61, p <0.0001, Fig.2a). Of the 139 genes found to show allele-specific expression a significant number are shared between reproductive and sterile workers (hypergeometric test p <0.0001, Fig.2b), with eight found only in sterile workers and 15 found only in reproductive workers (e.g. Fig.3, supplementary 1.0.1).

**Figure 2:**
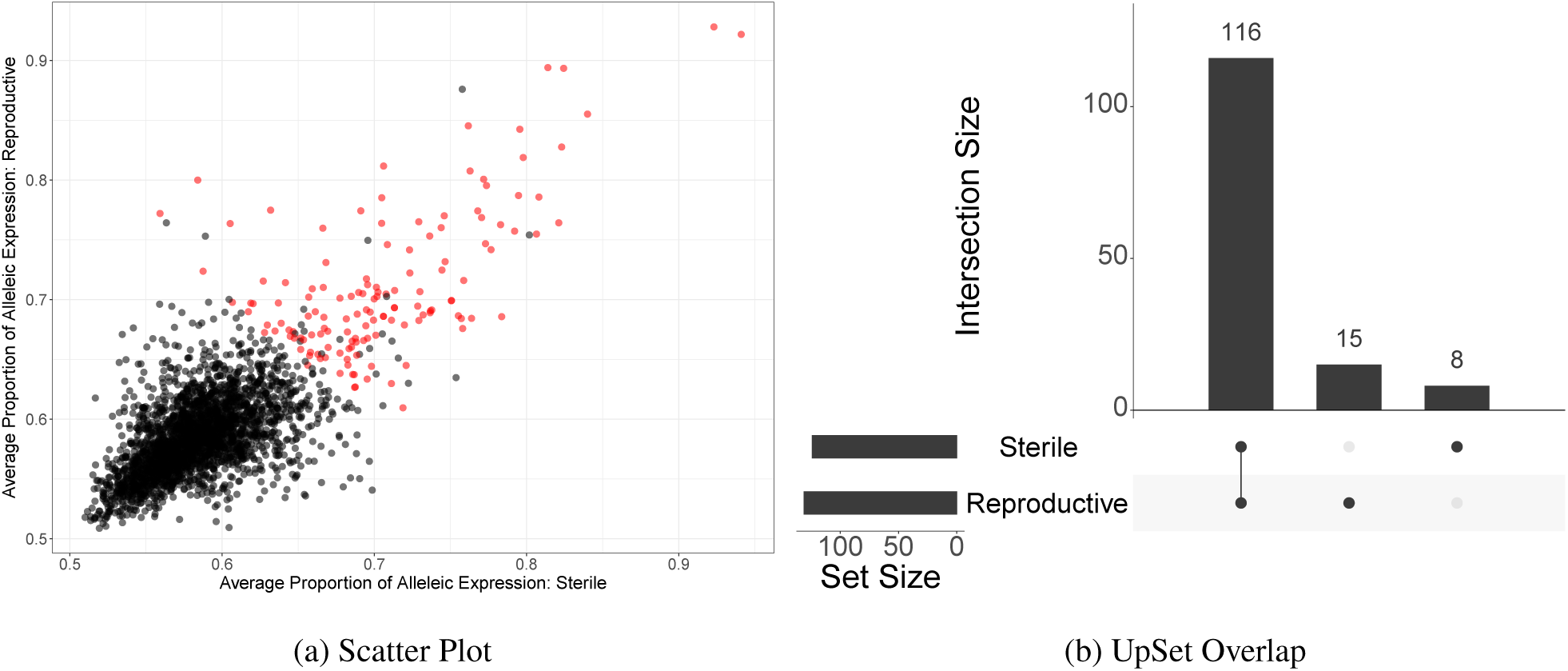
(a) Scatter plot showing the allelic expression proportion of sterile workers plotted against the allelic expression proportion of reproductive workers (allelic expression proportion averaged across colonies). Each point is a gene, the red points indicate genes showing significant allele-specific expression (q <0.05 and average allelic expression proportion >0.65). (b) An UpSet plot showing the number of allelically expressed genes shared by worker caste and the number unique to reproductive or sterile workers (intersection size), indicated by a joint dot or single dot respectively. The set size shows the total alleleically expressed genes in either reproductive or sterile workers.

**Figure 3:**
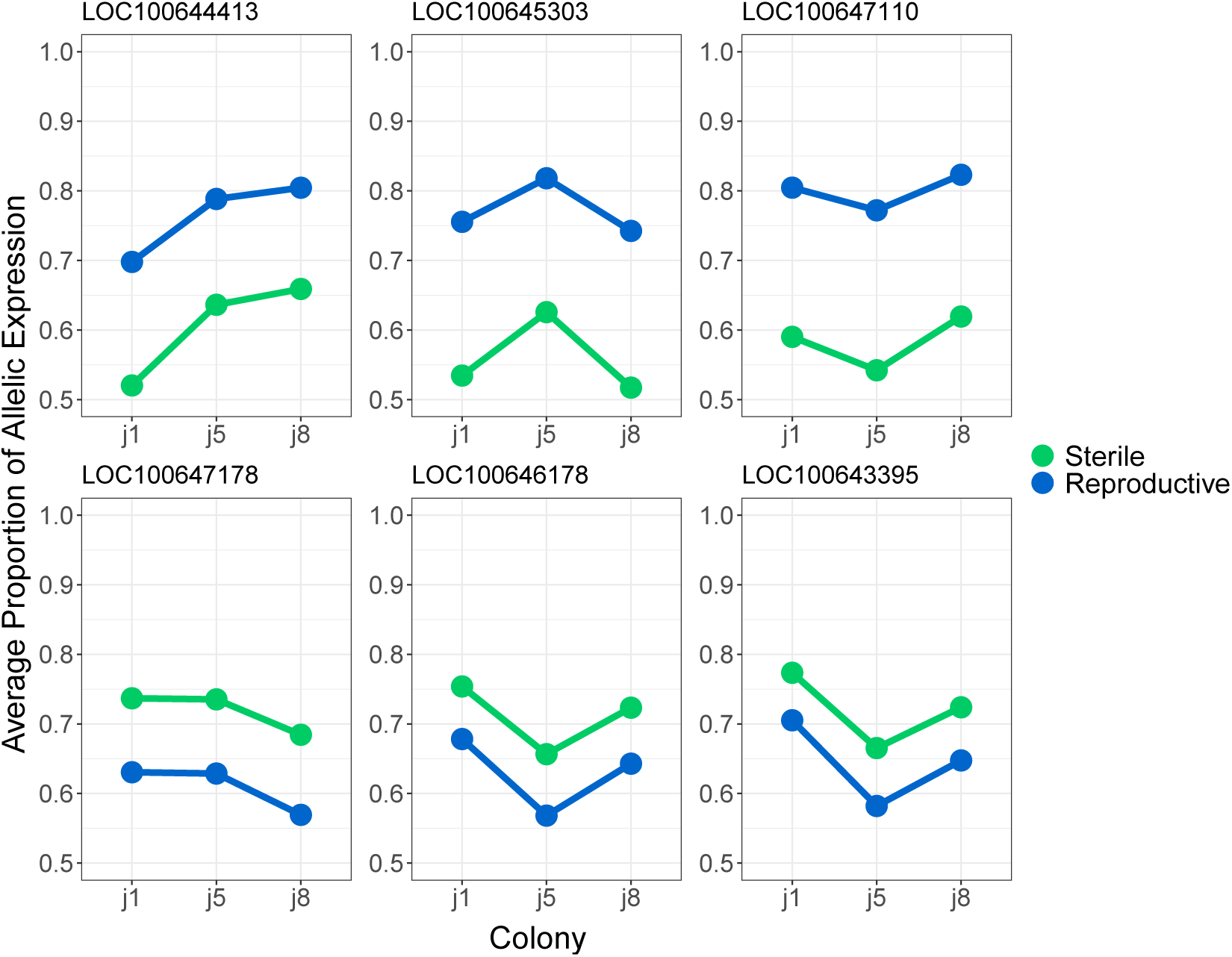
The average proportion of allelic expression for genes found to show significant allele specific expression in only sterile or reproductive workers across colonies. The top row shows the genes with the highest allelic expression bias in reproductive workers compared to sterile workers. The bottom row shows the highest allelic expression bias in sterile workers compared to reproductive workers.

There is also some variability in allelic expression proportion between colonies, with reproductive and sterile workers showing similar levels of bias compared to other colony replicates (supplementary 2.0 Fig.S5 and Fig.3). However, this is less apparent in the most highly biased genes (Fig.4).

**Figure 4:**
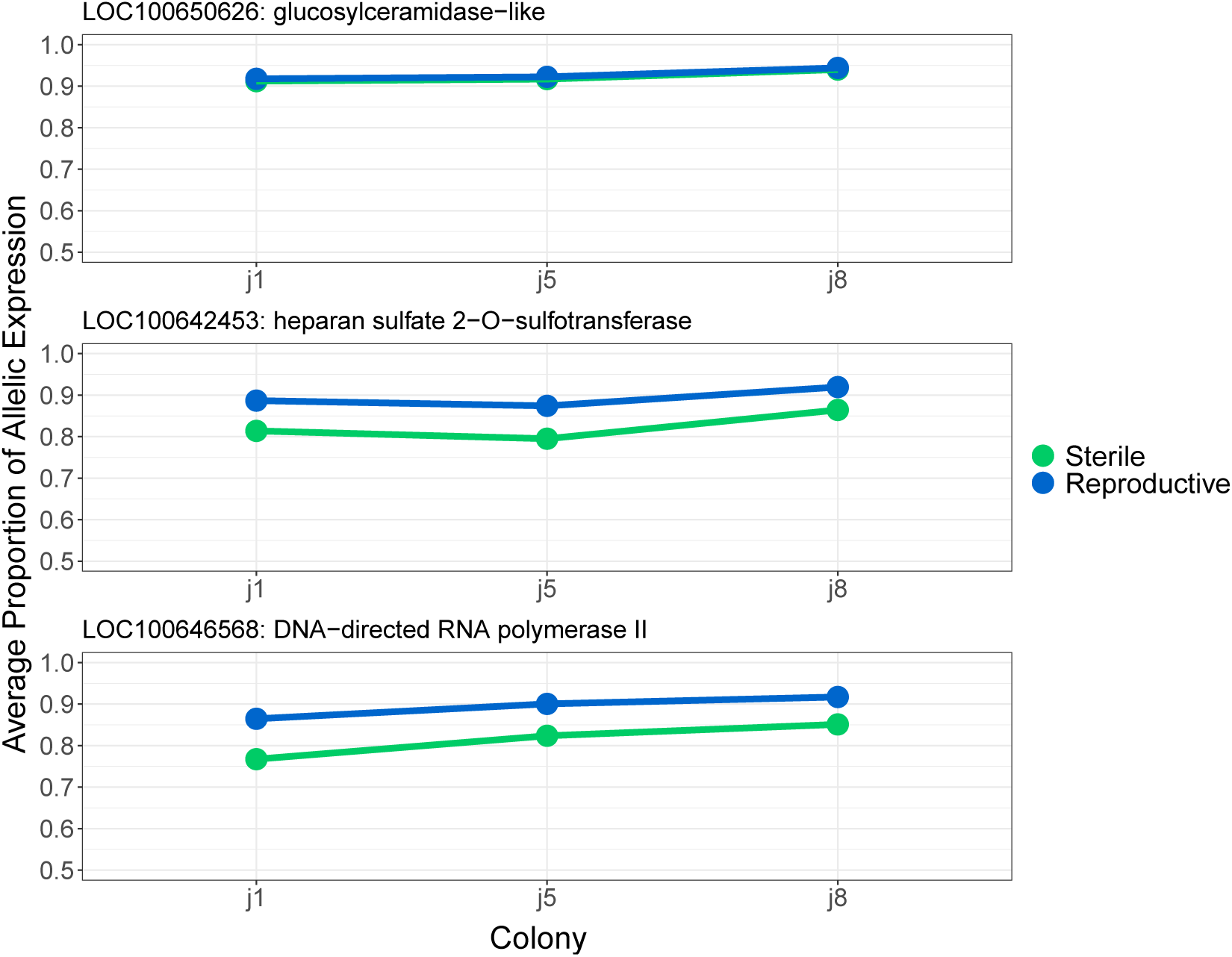
The average proportion of allelic expression for genes found to show the most extreme allele specific expression in both sterile and reproductive workers across colonies.

Enriched GO terms associated with genes showing significant allele-specific expression in both reproductive castes were involved in multiple biological processes, including; “*female gamete generation*” (GO:0007292), “*positive regulation of ovulation*” (GO:0060279) and “*histone H3-K27 acetylation*” (GO:0043974), supplementary 1.0.2.

GO terms enriched for the eight genes showing allele specific expression in sterile workers included mostly catabolic processes, but also “*response to pheromone*” (GO:0019236). The GO terms enriched for the 15 genes showing allele-specific expression in reproductive workers included; “*primary sex determination*” (GO:0007538) as well as multiple other cellular processes, supplementary 1.0.2. These results should be interpreted with care as the gene lists are relatively small. However, it is worth noting that the hypergeometric test used to generate the enriched terms has been previously shown to be the most appropriate statistic for gene ontology enrichment for small gene lists (Rivals *et al.*, 2007).

### Allele-specific methylation

Up to a maximum of 10bp were trimmed from the start of all reads due to base bias generated by the Illumina sequencing protocol (Krueger *et al.*, 2011). The mean mapping efficiency was 63.6% ± 1.4% (mean ± standard deviation) and the mean coverage was 17.7 ± 0.5 reads per base, the average number of uniquely mapped reads were 27,709,214 ± 753,203 (supplementary 1.0.3). 12.79% of the genome was not tested for allele-specific methylation as only regions in the main 18 linkage groups of the *B. terrestris* genome (Bter_1.0) could be tested.

Reproductive workers have significantly more allelically methylated regions compared to sterile workers, 303 (supplementary 1.0.4) compared to 201 (supplementary 1.0.5) respectively (Chi-squared goodness of fit; X-squared = 20.643, df = 1, p <0.0001). The majority of these regions occur within annotated genes, 26 and 15 allelically methylated regions occur outside of a gene for reproductive and sterile workers.

Most allelically methylated genes are unique to either sterile or reproductive workers, however, there is a significant number of common allelically methylated genes (hypergeometric test p <0.0001, Fig.5a). Most allelically methylated regions found within genes do not have additional annotation, however there are more located in exons compared to introns for both reproductive and sterile castes (Fig.5b).

**Figure 5:**
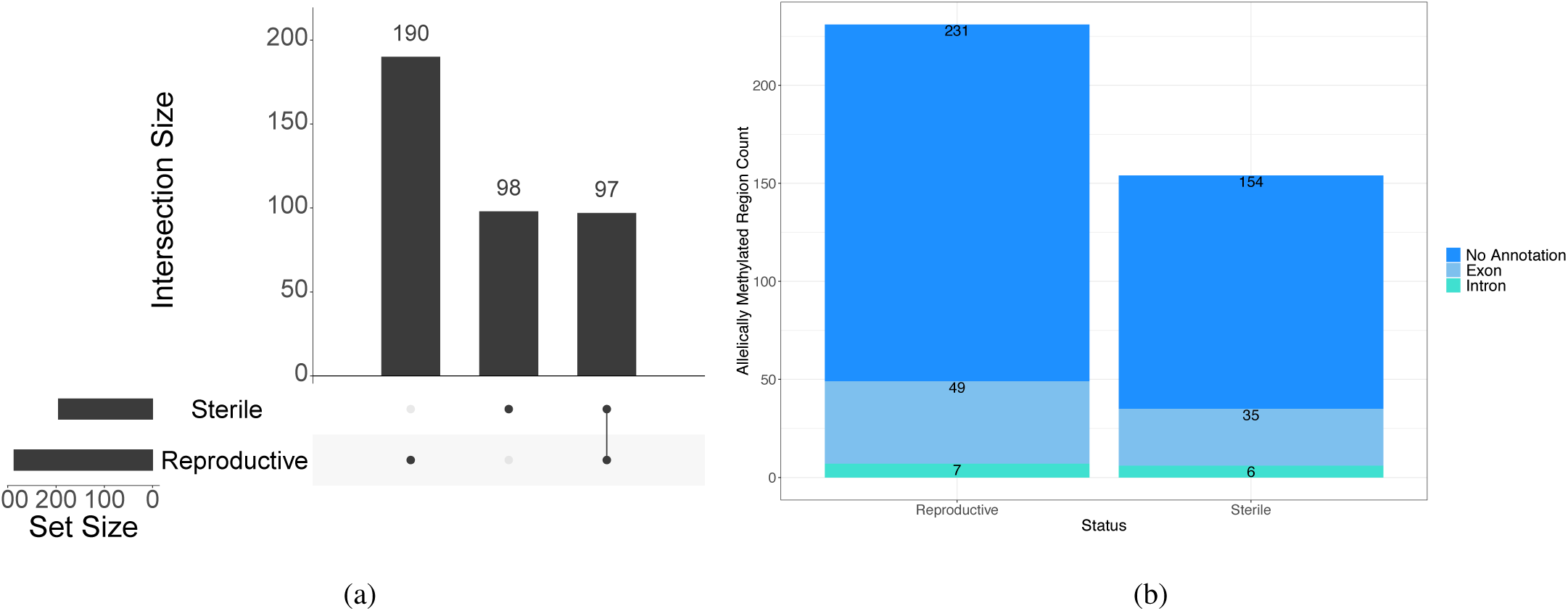
(a) UpSet plot showing the number of genes with allele-specific methylation in just reproductive and sterile workers, as well as the number of genes in common between both reproductive castes. (b) Component bar plot showing the number of allelically methylated regions within genes, found in exons and introns and the number without additional annotation.

Enriched GO terms associated with allelically methylated genes in both castes are involved in a variety of biological processes with many relating to the term “*positive regulation of RNA splicing*” (GO:0033120). As above, the enriched GO terms associated with allelically methylated genes in just sterile or reproductive workers are also involved in a large number of biological processes. However, the terms “*oocyte development*” (GO:0048599), “*ovarian follicle development*” (GO:0001541), “*oogenesis stage*” (GO:0022605) and other reproductive terms were enriched in allelically methylated genes of reproductive workers. Additionally none of these terms were identified in the GO terms associated with the allelically methylated genes of sterile workers (supplementary 1.0.6).

### Relationship of allele-specific expression and methylation

There is no significant overlap between genes showing allele-specific expression and allele-specific methylation (overlap between all conditions; hypergeometric test p = 0.209, Fig.6). However, six genes were found to show allele-specific methylation and expression in both reproductive castes, one gene was found to show allele-specific expression in both castes and allele-specific methylation in reproductive workers and one gene shows allele-specific expression in both castes and allele-specific methylation in sterile workers (Table 1).

**Figure 6:**
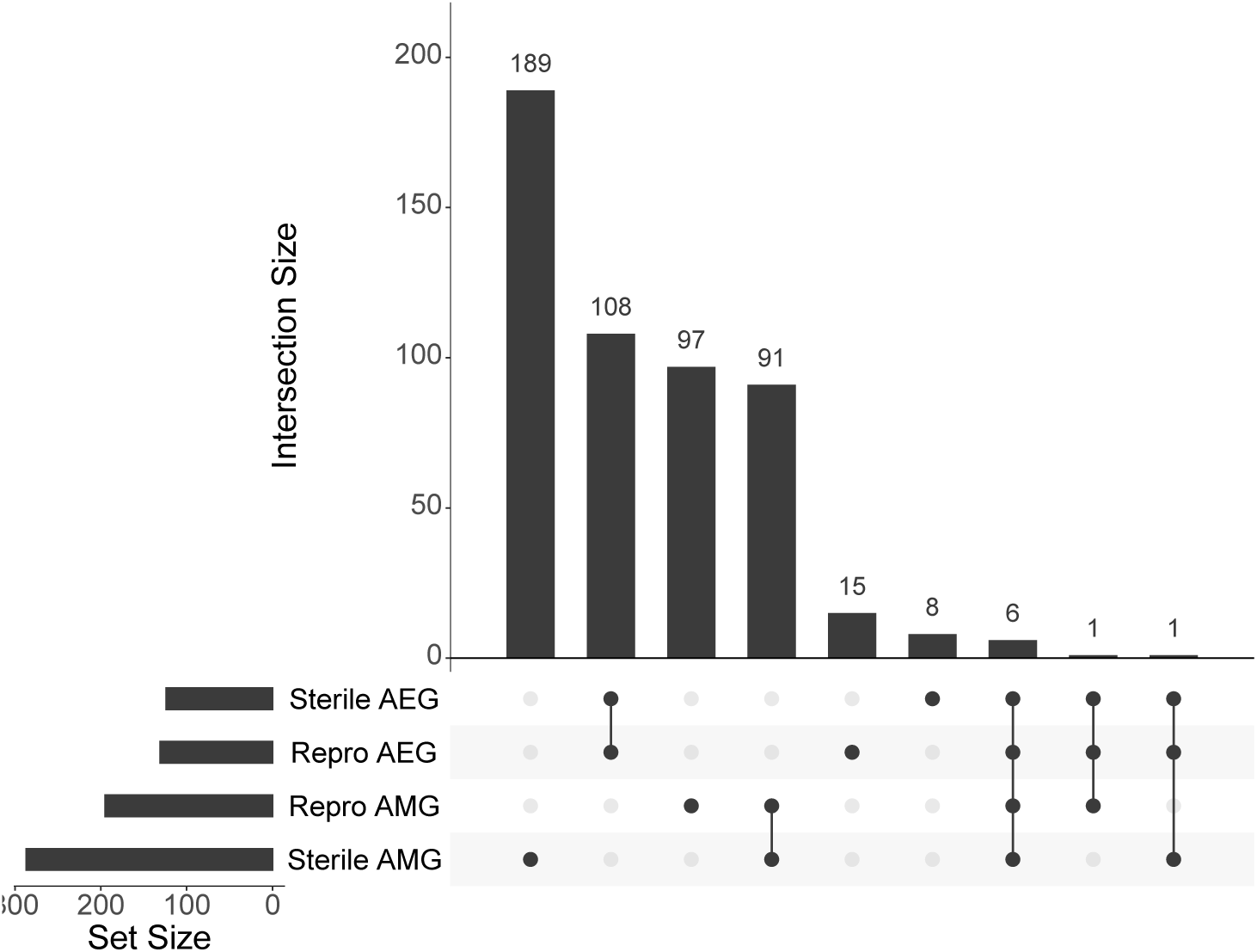
(a) UpSet plot showing the overlapping genes identified as allelically methylated and/or allelically expressed in both castes. AEG stands for allelically expressed gene. AMG stands for allelically methylated gene.

**Table 1:** Genes identified as showing allele-specific methylation and expression in both reproductive castes. * this gene does not show allele-specific methylation in sterile workers. ^◊^ this gene does not show allele-specific methylation in reproductive workers.

| Gene ID | Gene Description |
| --- | --- |
| LOC100643777 | 40S ribosomal protein S6 |
| LOC100643941 | connectin |
| LOC100644811 | neuroligin-4, Y-linked |
| LOC100652132 | importin-11 |
| LOC100644932 | AP-1 complex subunit mu-1 |
| LOC105665778 | regulator of microtubule dynamics protein 1-like |
| LOC105666711* | tyrosine-protein kinase Btk29A* |
| LOC100643219° | putative pre-mRNA-splicing factor ATP-dependent RNA helicase PRP° |

The GO terms enriched for the genes found to be allelically methylated and expressed (Table 1) compared to the entire genome as background, included a large variety of biological processes (supplementary 1.0.7). Specifically some reproductive related terms were enriched; “*female germline ring canal formation*” (GO:0007301) and “*ovarian fusome organization*” (GO:0030723).

There is a significant difference in the proportion of allelic expression of genes allelically methylated in either reproductive workers, sterile workers or both (Kruskal-Wallis; chi-squared = 28.838, df = 2, p <0.0001). Genes allelically methylated in both castes show on average higher levels of allele specific expression compared to those unique to either reproductive or sterile workers (Dunn test with Benjamin-Hochberg correction; both compared to unique in reproductive workers Z = 5.149, q <0.0001, both compared to unique in sterile workers Z = 4.147, q <0.0001), Fig.7. Additionally, genes with allele-specific methylation unique to reproductive workers show similar levels of allelic expression compared to genes with allele-specific methylation unique to sterile workers (Dunn test with Benjamin-Hochberg correction; reproductive compared to sterile Z = −1.851, q = 0.06), Fig.7. Finally, there is no interaction between reproductive caste and allelic expression proportion on the allelic-methylation status of a gene (Anova, interaction vs main effects model, F_2, 296_ = 0.1094, p = 0.896), Fig.7.

**Figure 7:**
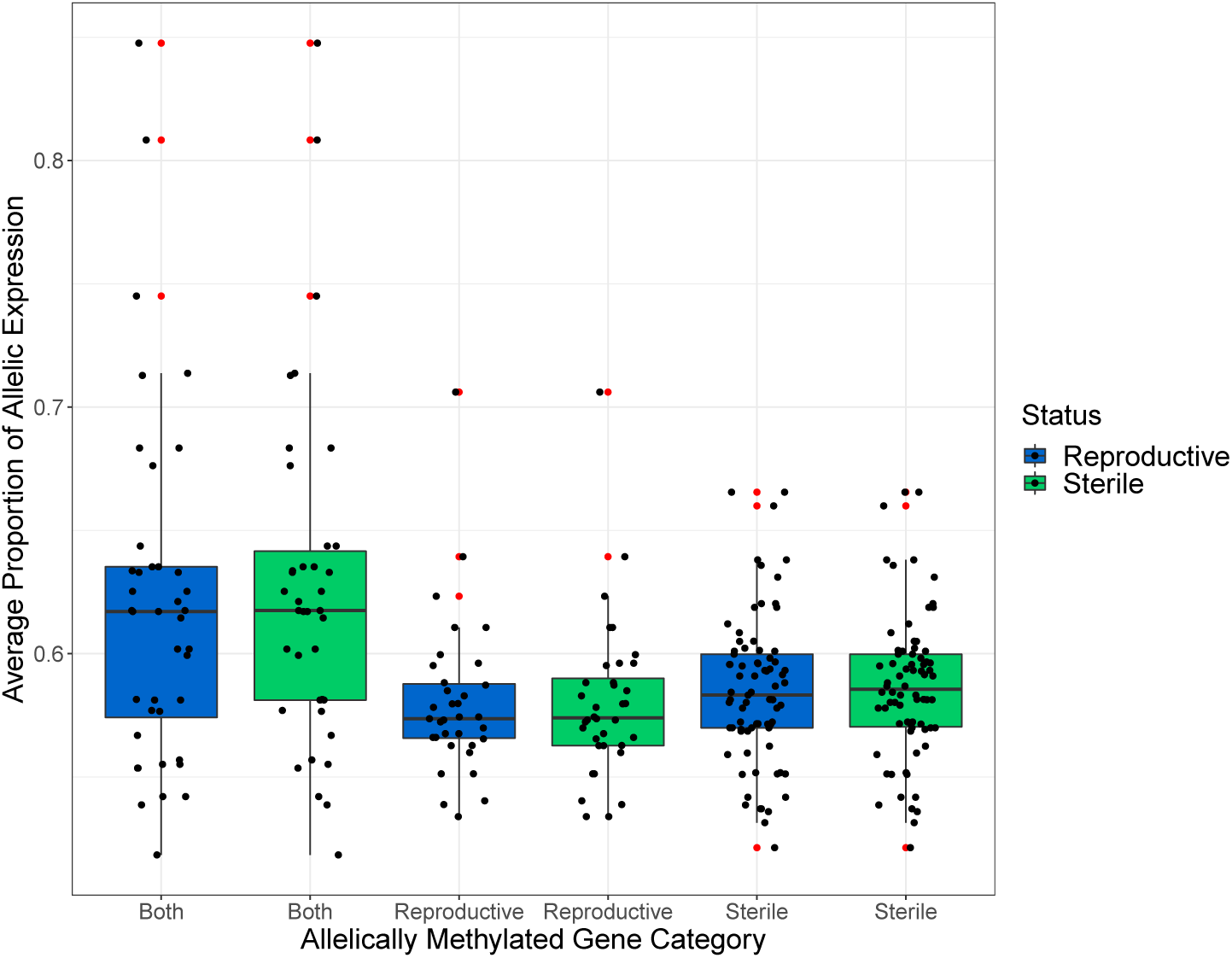
Boxplots showing the proportion of allelic expression in reproductive and sterile workers for genes identified as allelically methylated in either both castes, just reproductive workers or just sterile workers. Each boxplot shows the median along with the 25th and 75th percentile. The whiskers represent 1.5X the interquartile range. Outliers are represented as additional red points and each gene is represented by a black dot.

## Discussion

Using whole genome bisulfite sequencing and RNA-seq from reproductive and sterile *B. terrestris* workers from three independent colonies we have identified genome wide allele-specific expression and allele-specific methylation. The majority of genes displaying allele-specific expression are common between reproductive castes and the proportion of allele-specific expression generally varies between colonies. This study has also identified allele-specific methylation differences between reproductive castes and found the majority of allelic-methylation events are located within genes. However, there is no significant overlap of genes showing allele-specific expression and allele-specific methylation. We have also found that genes with common allele-specific methylation between castes show a higher proportion of allelic-expression bias compared to allelically methylated genes unique to either reproductive or sterile workers.

This study has identified 139 genes which show allele-specific expression from a stringent subset of genes covering 24% of all annotated genes within the *B. terrestris* genome. This number is in line with previous research that identified around 500 loci across the whole genome of *B. terrestris* (Lonsdale *et al.*, 2017). Predictions based on the kinship theory suggest if imprinted genes exist in social insects, such as *B. terrestris*, they will be involved in worker reproductive behaviour (Queller, 2003). GO terms enriched for the allelically expressed genes were involved in reproduction amongst other biological processes. This finding supports the idea that if imprinted genes are present in *B. terrestris* some will be involved in worker reproductive behaviour.

The proportion of allelic expression bias differed between colonies and the GO terms enriched for all allelically expressed genes, whilst containing reproductive terms, were varied. This indicates allele-specific expression may be involved in other mechanisms, rather than solely imprinting, and that it plays a diverse role in *B. terrestris*. Previous research identified 61 genes showing allele-specific expression in a cross of two *Nasonia* species, the expression bias in all genes was attributed to *cis*-effects (Wang *et al.*, 2016). There have also been a number of non-imprinted loci found in humans which show allele-specific expression directly associated with *cis*-acting polymorphic sites, such a single nucleotide polymorphisms (SNPs) (Tycko, 2010). Given that each colony used here is genetically distinct, *cis*-effects, such as SNPs, are likely represented in the results. In humans <1% of genes are imprinted but considerably more exhibit allele-specific expression (Tycko, 2010), it is therefore reasonable to assume only a small percentage of the genes identified as showing allele-specific expression in this study may actually be imprinted genes.

Whilst the majority of genes showing allele-specific expression were common between reproductive castes, a large number of genes show allele-specific methylation which is unique to either reproductive or sterile workers. Additionally, there are significantly more allelically-methylated sites in reproductive workers compared to sterile workers, with allelically methylated genes in reproductive workers enriched for GO terms related to reproduction. These findings support previous research which suggests methylation is associated with worker reproductive behaviour. Amarasinghe *et al.* (2014) found a global erasure of DNA methylation increased reproductive behaviour, Liu *et al.* (2018) found differences in expression in genes responsible for methylation between castes and Marshall *et al.* (2019) found differentially methylated genes between *B. terrestris* castes, some of which were involved in reproductive processes. Numerous other studies have linked methylation to caste differences in various other social insect species, such as; *Apis mellifera* (Lyko *et al.*, 2010; Elango *et al.*, 2009), *Camponotus floridanus* and *Harpegnathos saltator* (Bonasio *et al.*, 2012), *Polistes dominula* (Weiner *et al.*, 2013) and *Zootermopsis nevadensis* (Glastad *et al.*, 2016). The development of experimental techniques to alter DNA methylation, such as CRISPR/Cas (Vojta *et al.*, 2016), will allow for experiments to test the causal effect of DNA methylation and allele-specific methylation on caste determination in social insects.

It is, however, clear from this study that DNA methylation does not play a direct causal role in the production of all allele-specific expression events, with only a small number of genes displaying both allele-specific expression and methylation. This does not rule out the possibility that methylation may act as an imprinting mark, if only a small number of genes are actually imprinted, as in humans (Tycko, 2010). GO terms enriched for the few genes which do show allele-specific methylation and allele-specific expression included some reproductive related terms. As the kinship theory predicts imprinted genes should affect reproduction in *B. terrestris* (Queller, 2003), the identification of these genes provides the groundwork for future research to further investigate the possibility of parent-of-origin methylation as an imprinting mark.

Additional imprinting marks should not be ruled out however as GO terms enriched for genes showing allele-specific methylation included histone modifications. Genes displaying allele-specific methylation may feed into other mechanisms which may, in-turn, drive allele-specific expression, accounting for the lack of direct association. For example, methylation of an imprinting control region can signal certain histone modifications which can allow the formation of condensed chromatin, silencing many genes in one region (Barlow, 2011), this process can also occur in an allele-specific manner (Tycko, 2010).

Whilst only a small number of genes show allele-specific methylation and allele-specific expression, genes showing allele-specific methylation in both reproductive castes had higher allelic expression bias compared to those found only in one caste. One explanation is that allelically methylated genes present in both castes carry out different functions to those identified in a single caste. This is supported by the diverse GO terms obtained for shared and caste-specific allelically methylated genes. In humans, the majority of allele-specific methylation is genotype dependent rather than parentally inherited (Meaburn *et al.*, 2010). Whereas, allele-specific methylation associated with imprinting may change at different stages of development (Edwards *et al.*, 2017). It may therefore be that the common allelically methylated genes identified here are linked to genotype (i.e. epialleles) whereas the caste-specific allelically methylated genes may represent imprinting marks. However, this is speculation and requires further investigation.

In order to further understand the role and origin of allele-specific methylation a pipeline is needed which integrates SNP data (generated from genomic DNA), to allow the identification of specific alleles. Using this method rather than the probabilistic models employed here would enable hyper/hypomethylation (i.e. higher or lower methylation in one conditions compared to another) to be associated with allele-specific expression when they occur in tandem. Additionally, this method, with increased biological replication per colony, would facilitate the identification of epialleles, i.e. when allele-specific methylation is driven by genotype. Epialleles have been identified in the honeybee (Wedd *et al.*, 2016) and will be important in the identification of parent-of-origin methylation (Remnant *et al.*, 2016).

Overall, this study provides evidence suggesting imprinted genes exist in *B. terrestris* and are related to worker reproductive behaviour as predicted by Queller (2003) and Haig (2000). However, the role of DNA methylation, as a mechanism of potential imprinting, is still unclear. The diverse function of genes showing allele-specific expression and/or allele-specific methylation suggests a varied role for these genomic mechanisms and experimental validation of their function is required. Future research utilising highly related reciprocal crosses, in order to identify the parental origin of an allele, is needed to discover imprinted genes and to take into account *cis*-effects, such as genotype. These types of crosses can also be used to further investigate parent-of-origin DNA methylation as a mechanism of imprinting.

## Acknowledgements

This research used the ALICE2 High Performance Computing Facility at the University of Leicester. H.M. was supported by a NERC CENTA DTP studentship. A.R.C.J. and Z.N.L. were supported by BBSRC MIBTP DTP studentships. This work was supported by the Natural Environment Research Council [grant number: NE/N010019/1 to E.B.M.]. The authors declare no conflict of interests.

## Author contributions

E.B.M. conceived the study. H.M. analysed the data. A.R.C.J. and Z.N.L. contributed to the allele-specific expression analyses. H.M wrote the initial manuscript. All authors contributed to and reviewed the manuscript.

## Data Accessibility

Data has been deposited in GenBank under NCBI BioProject: PRJNA533306. All code is avaliable at http://doi.org/10.5281/zenodo.1974852.

