## Supplementary_2.0 for "Bumblebee worker castes show differences in allele-specific DNA methylation and allele-specific expression"

### 2.0 Additional figures

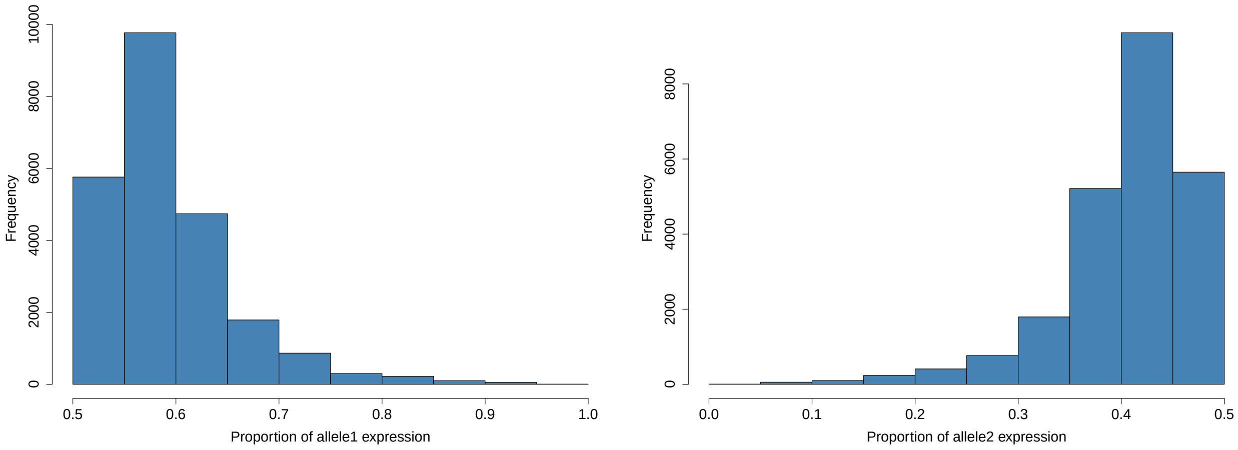


(a) Allele 1 (b) Allele 2

Figure S1: Histograms of the average forced expression proportion across all SNPs and samples for the hypothetical alleles one and two.

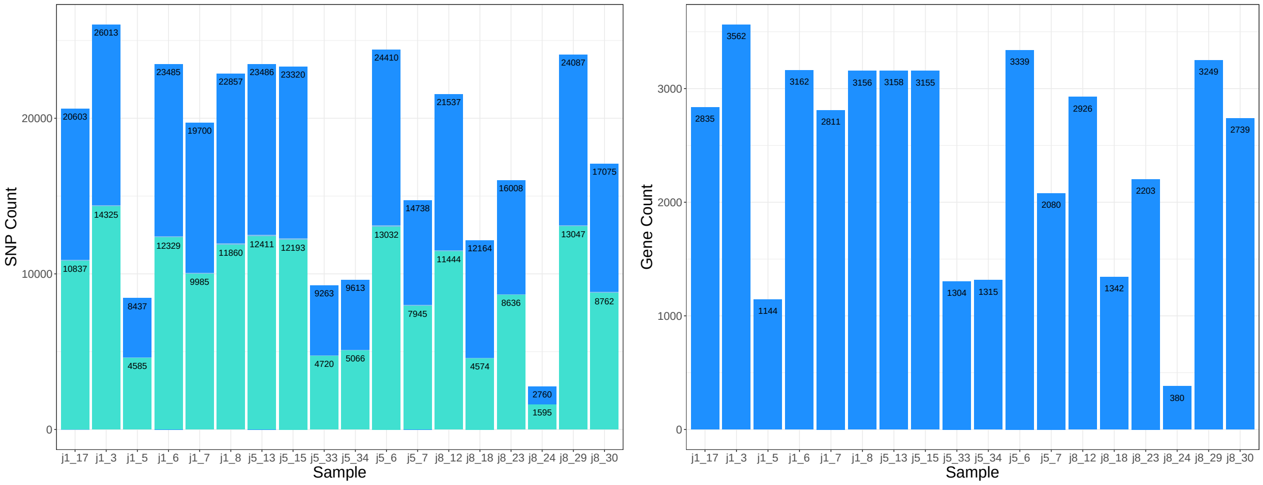


(a) SNP Counts (b) Gene Counts

Figure S2: (a) Component bar chart showing the total number of SNPs called per sample and the number of those which have a minimum coverage of 10 and coverage for both alternate and reference alleles. (b) The number of genes per sample which have at least one SNP present. The sample names represent the unique colony (j1, j5 or j8) and the individual number.

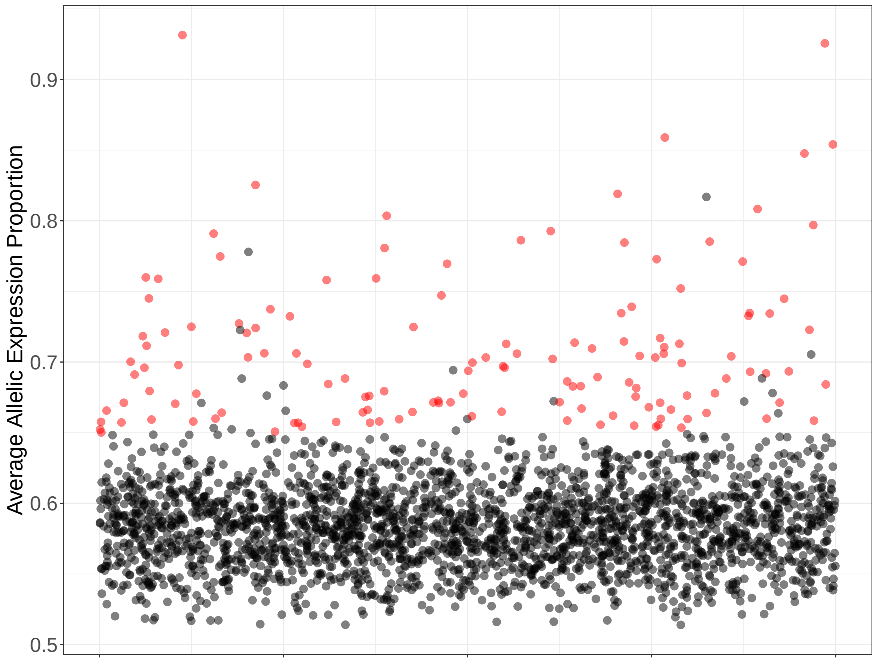


Figure S3: Spread of allelic ratios averaged across all colonies and reproductive samples. Each dot represents a gene, the red dots show genes showing significant allelic expression bias (q <0.05 and average allelic expression proportion >0.65).

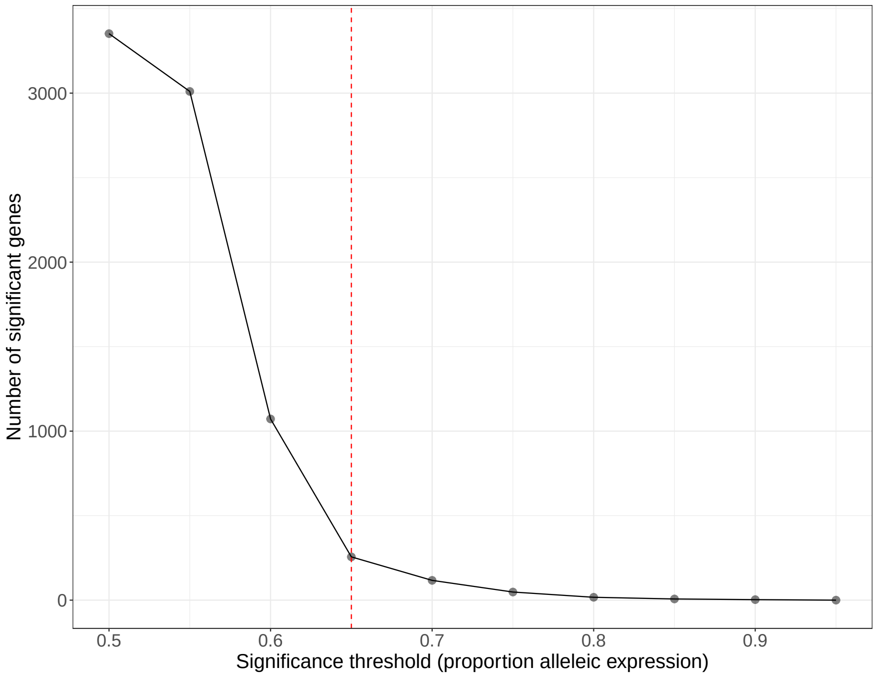


Figure S4: Plot showing the number of genes which are allelically expressed (q <0.05) per arbitrary cut-off expression proportion. The red dashed line indicates the cut-off threshold chosen of 0.65.

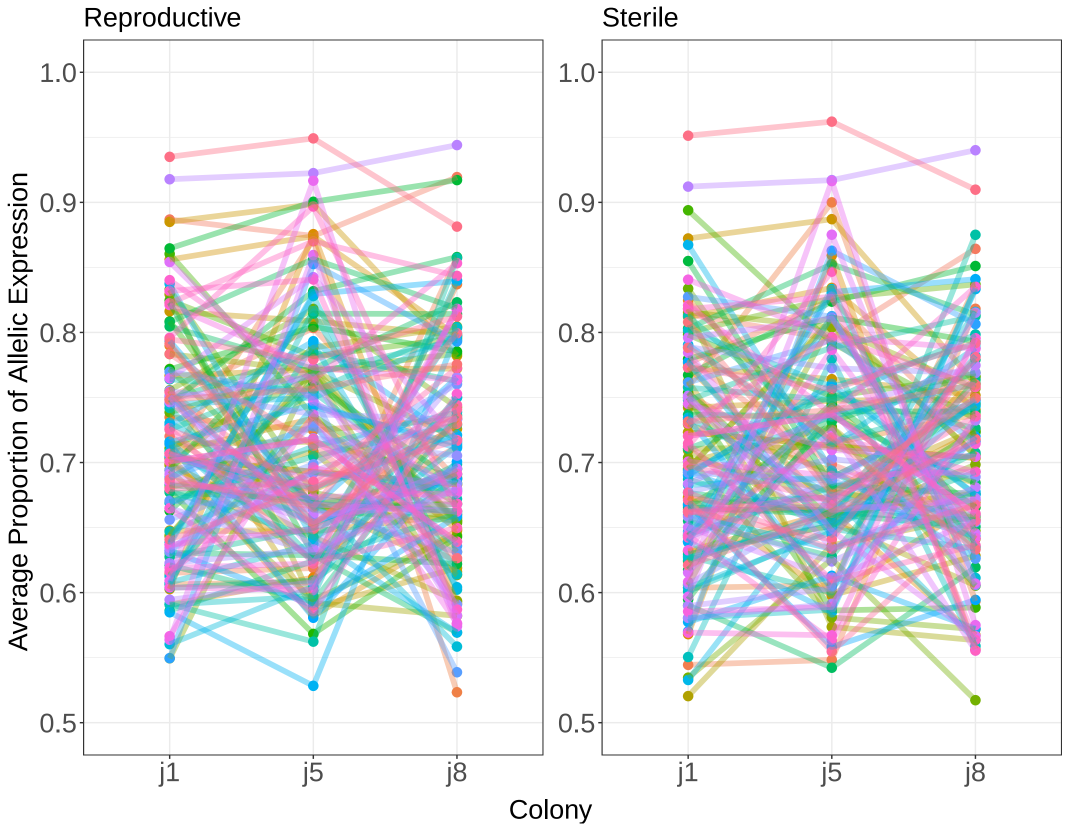


Figure S5: The average proportion of allelic expression for reproductive and sterile workers across each colony. Each colour represents a unique gene. Genes shown are those which show significant allele-specific expression (q <0.05 and average proportion of expression bias >0.65).
